# Uncovering a new player in ischemic stroke: a study of intra-arterial interferon-gamma-producing monocytes in hyperacute stroke

**DOI:** 10.1101/2025.08.19.671149

**Authors:** Katherine Hernandez, Erik J. Plautz, Safia Sharif, Nathan Jones, Nneka Osiah, Sterling B. Ortega

**Affiliations:** Department of Microbiology, Immunology, and Genetics, University of North Texas Health Science Center, Fort Worth, TX; Department of Neurology, University of Texas Southwestern Medical Center, Dallas, TX; Texas College of Osteopathic Medicine, University of North Texas Health Science Center, Fort Worth, TX; College of Natural and Health Sciences, Virginia State University, Petersburg, VA

**Keywords:** monocytes, interferon-gamma, stroke, tMCAO, cytokines, inflammation, CD14

## Abstract

Stroke triggers a rapid and complex immune response that is not yet fully understood, especially within hours after an ischemic infarct. Our previous study in stroke patients revealed a significant increase in interferon- gamma (IFN-γ) immediately (hyperacute) and downstream of the ischemic ictus, within the arterial compartment. The present study investigated the source, inciting factors, and role of IFN-γ in a preclinical murine model. Stroke was produced using transient middle cerebral artery occlusion, and immune cells within the arterial vasculature distal to the occlusion (pre- and post-occlusion) were characterized using flow cytometry. Compared with the control samples, the post-occlusion samples presented an increase in IFN-γ^+^ and CD69^+^ cells, whereas no significant increase was detected in IL17^+^, IL4^+^, and CD25^+^ cells. Further analysis of the IFN-γ^+^ population revealed two novel attributes. First, interrogation of the identity of these IFN-γ^+^ cells revealed that the increase in IFN-γ production was largely driven by CD14^+^ cells in the post- occlusion sample, with negligible contributions from other canonical IFN-γ-producing cells (CD4, CD8). Second, the IFN-γ^+^ cells exhibited two distinct clusters, an IFN-γ^low^ and an IFN-γ^hi^ population. Further analysis revealed that the IFN-γ ^low^ population was largely composed of CD14^+^ cells, whereas the IFN-γ^hi^ population was dominated by CD4^+^ T-cells. To explore the conditions driving IFN-γ production, an *in vitro* ischemia model involving oxygen-glucose deprivation (OGD) was employed. Co-culturing of naïve splenocytes with OGD-treated CNS cells and OGD-derived supernatant resulted in a significant increase in IFN-γ^+^CD14^+^ cells, as compared to normoxic controls, an effect that coincided with marked loss of DAPI^+^ and NeuN^+^DAPI^+^ cells in mixed cortical (neuronal and glial) cultures. In summary, this study identified intra-arterial CD14^+^ monocytes as novel early sources of IFN-γ in the hyperacute phase of stroke, a role traditionally attributed to adaptive immune cells. Using *in vivo* and *in vitro* ischemia models, the findings reveal that injury-associated signals from CNS cells are sufficient to directly induce IFN-γ production in CD14^+^ cells, redefining early stroke immunopathology and uncovering a potential target for timely immunomodulation.

## INTRODUCTION

Stroke is a leading cause of long-term disability and death worldwide, with over 12.2 million new cases and 6.6 million deaths annually [1]. It is caused by the sudden interruption of blood flow in the central nervous system (CNS), leading to rapid neuronal injury and the initiation of a complex sequence of neurovascular and immune responses that evolve over time and are still under considerable research. Seminal work elucidating the acute (days) and chronic (months) phases of stroke has furthered our understanding of the pathophysiology of stroke [2,3]. However, studies focused on the first few hours after a stroke, referred to as the hyperacute phase, remain poorly characterized. During this critical window, molecular and cellular changes set the stage for subsequent tissue injury, repair, and recovery.

The current standard of care for ischemic stroke patients includes intravenous thrombolysis with tissue plasminogen activator (tPA) within 4.5 hours of onset or endovascular thrombectomy for up to 16-24 hours in select patients [4–7]. Moreover, the ictus initiates a rapid and coordinated immune response characterized by the activation of CNS-resident microglia and shifts from a surveillance to an activated phenotype, releasing pro-inflammatory cytokines, including TNF-α, IL-1β, and IL-6. Once infiltration into the CNS is facilitated, neutrophils appear within 6–24 hours, releasing reactive oxygen species (ROS) and proteolytic enzymes such as matrix metalloproteinase-9 (MMP9), which further amplify local CNS inflammation and facilitate additional peripheral immune infiltration. Monocyte infiltration occurs more gradually, with the earliest infiltration occurring within 24 hours and the peak occurring 2-3 days post-stroke [8].

Monocytes are circulating immune cells of the innate immune system that are produced in the bone marrow and found in the blood. They are characterized by their expression of CD14, the coreceptor for lipopolysaccharide (LPS). Monocytes can be further classified into functional subsets on the basis of surface marker expression: classical monocytes are proinflammatory, rapidly recruited into inflamed tissues, and are characterized by their robust expression of Ly6C and lack of CD16 expression; nonclassical monocytes are involved in patrolling and tissue repair and are characterized by low expression of Ly6C and robust expression of CD16. In line with this dichotomy, studies focused on the acute and chronic phases after stroke have revealed a context-dependent role that spans both the pathogenic and reparative spectra. During the acute phase, classical monocytes from the bone and spleen are recruited to the CNS by CCL2, infiltrating the brain tissue within 24–72 hours [9]. In the brain parenchyma, the monocytes differentiate into macrophages and adopt a predominantly proinflammatory phenotype, secreting cytokines such as TNF-α, IL-1β, and IL-6, producing nitric oxide and ROS, and promoting blood–brain barrier (BBB) disruption [10]. At 3–7 days post-stroke, monocyte-derived macrophages shift toward a reparative phenotype, producing IL-10, TGF-β, and growth factors that support angiogenesis, oligodendrocyte survival, and axonal sprouting [11,12]. During the chronic phase (months), it has been shown that monocytes are involved in the clearance of cellular debris, remyelination, and the modulation of glial scar formation [11,12]. However, the events during the hyperacute phase, specifically the primordial immune events occurring just after ischemia and within the arterial vasculature, remain poorly understood. Our prior research uncovered notable findings in stroke patients: during the hyperacute phase (7 hours from the last known well time) of stroke onset, the arterial post-ictus environment presented elevated levels of MMP9 and IFN-γ [13]. This observation was perplexing, as it exhibited a temporal anomaly characterized by the production of IFN-γ so soon (hours) after ischemia, and a spatial anomaly characterized by the production of IFN-γ within the occluded arterial vasculature, warranting further study.

IFN-γ is a pleiotropic cytokine traditionally associated with an active adaptive immune response. Produced primarily by natural killer cells, helper type-1 T-cells, and cytotoxic type-1 T-cells, IFN-γ plays a central role in enhancing antigen presentation and promoting macrophage activation [14]. The role of IFN-γ in brain health remains controversial, with studies indicating both neuropathogenic and neuroprotective effects, depending on factors such as the cellular source, target, timing, and concentration [15–18]. Recent investigations, however, have revealed the production of IFN-γ by other adaptive immune cells, such as B-cells [19–21]. Although IFN- γ is not typically associated with innate cells, increasing evidence suggests that dendritic cells [22,23], monocytes [24], and macrophages [25,26] can produce IFN-γ. Notably, monocytes are generally not primary producers of IFN-γ; rather, they are more commonly targets of IFN-γ signaling [27].

Using a combination of *in vivo* and *in vitro* models, as well as cellular and microscopic approaches, we expanded upon this discovery and delved deeper into the cellular origins, role, and inciting insult. In this work, we recapitulate the temporal and spatial anomalies by showing an increase in IFN-γ-producing cells within the arterial post-occlusion environment within hours of stroke induction in an animal model. Using a murine stroke model in combination with flow cytometry, *in vitro* OGD, and immune-CNS co-culture systems, we identified post-ictus intra-arterial CD14^+^ monocytes as rapid and context-dependent sources of IFN-γ, characterized their activation dynamics, and uncovered conditions under which they may contribute to early neuroinflammatory injury. These results reveal a novel role for innate myeloid cells in shaping the hyperacute immune landscape of stroke and provide insight into an underexplored window of therapeutic opportunity.

## MATERIALS AND METHODS

### Mice

Male and female C57BL/6J mice (9–12 weeks old) were obtained from The Jackson Laboratory (Bar Harbor, ME) and housed in a pathogen-free facility at the University of North Texas Health Science Center (UNTHSC). The mice were group-housed (4–5 per cage) in standard animal housing with corncob bedding and environmental enrichment. Cages were randomly assigned to include multiple experimental groups. All animals were maintained on a 12-hour light/dark cycle with *ad libitum* access to standard rodent chow and autoclaved water. All experimental procedures were approved by the Institutional Animal Care and Use Committee (IACUC) at the UNTHSC (Protocol # 2021-0045).

### Stroke Model

Transient middle cerebral artery occlusion (tMCAO) was performed on C57BL/6J mice with 2% isoflurane in a 70% nitrous oxide and 30% oxygen mixture [28]. The mice were placed on a heating pad to preserve normothermia throughout the procedure and anesthesia depth was confirmed via a lack of withdrawal reflex to toe pinch. After surgical exposure, the carotid artery was ligated, and a 0.23 mm nylon monofilament (Cat. 602334PK10, Doccol Corporation, Sharon, MA) was introduced into the internal carotid artery and advanced to the middle cerebral artery (MCA) to induce occlusion for 60 minutes. Successful occlusion was confirmed by a ≥ 80% reduction in cerebral blood flow compared with that at baseline, as measured using a laser Doppler flowmeter (Moor Instruments Inc., DE). Afterwards, the filament was withdrawn to allow reperfusion; restoration was considered successful with ≥ 50% recovery of baseline blood flow. Immediately following surgery, 0.25% bupivacaine was administered subcutaneously at the incision site. The animals were monitored during recovery and provided moistened chow and saline as needed.

### Blood Collection

The pre-occlusion blood samples were collected from the carotid artery after incision (but immediately before filament insertion) and placed into an acid-citrate dextrose (ACD, yellow top) tube (Cat. 364606, BD Biosciences, NJ). The post-occlusion blood sample was collected immediately after suture retraction but before full reperfusion. The distal control sample was collected via submandibular blood draw from a separate cohort of naïve animals not subjected to tMCAO, to avoid confounding inflammation from surgery. For this procedure, the mice were gently restrained, the area was sterilized with an alcohol wipe, and the vessel was punctured with a sterile lancet; blood was collected into ACD tubes.

### Tissue Sample Preparation

Spleens were harvested and processed into single-cell suspensions by mechanical dissociation through a 70 µm nylon filter (Corning, NY), followed by rinsing with complete culture media. The complete culture media consisted of 1X Roswell Park Memorial Institute (RPMI, Cat. MT10040CV, Corning, NY) supplemented with 10% fetal bovine serum (FBS, Cat. 100-106-50, Gemini Bio, CA), 1.25% HEPES buffer (Cat. 15630080, Gibco, NY), 1% non-essential amino acids (Cat. 11140050, Gibco, NY), 1% penicillin-streptomycin (Cat. 15140122, Gibco, NY), 1% L-glutamine (Cat. 25030081, Gibco, NY), and 0.0002% 2-mercaptoethanol (Cat. M3148-25ML, Sigma, JP). The resulting single-cell suspension was subsequently centrifuged. After centrifugation, the supernatants were discarded, and the pellet was resuspended in 5 mL of fresh culture media. To remove debris, red blood cells, and dead cells, Lympholyte-M (Cat. CL5035, Cedarlane Labs, CAN) was used according to the manufacturer’s instructions. Briefly, Lympholyte-M was added to a 15 mL conical tube, and the splenocyte suspension was gently layered on top [29]. The sample was subsequently centrifuged at 1000 × g for 20 minutes at 21°C without braking. The resulting buffy coat was collected and transferred to a new conical tube, followed by resuspension in culture medium and a second centrifugation. After the supernatant was discarded, the pellet was resuspended in fresh culture medium. Cell viability and counts were assessed via the use of 4% trypan blue (Cat. 15250061, Gibco, NY) and a Cytosmart cell counter (Corning).

### CD14^+^ Cell Isolation

CD14^+^ cells were isolated using the EasySep Mouse PE Positive Selection Kit II (Cat. 17684, Stemcell, CAN) following the manufacturer’s instructions. A single-cell suspension of splenocytes was resuspended in RoboSep Buffer (Cat. 17684RF, Stemcell, CAN), and Fc receptor (FcR) blocking reagent was added to the suspension. An anti-mouse CD14 PE-conjugated antibody (Cat. 150105, BioLegend, CA) was added, and the cells were incubated at room temperature for 15 minutes. After incubation, the cells were washed with RoboSep Buffer and centrifuged at 300 × g for 5 minutes at room temperature. The supernatant was discarded, and the pellet was resuspended in the original volume of RoboSep Buffer. The selection cocktail was added, and the sample was incubated at room temperature for 15 minutes. Rapid Spheres were added, and the volume was adjusted to 2.5 ml with RoboSep Buffer. The mixture was incubated at room temperature for 5 minutes. The supernatant was aspirated and discarded. This wash step was repeated three additional times, for a total of four incubations. Following the final wash, the CD14^+^ cells were resuspended in complete culture medium for downstream *in vitro experi*ments.

### Flow Cytometry Analysis

Following blood collection or post-culture incubation, cellularity was quantified using fluorescence-activated cell sorting (FACS) analysis. Survey assay was performed as previously described but tailored for blood isolation during the tMCAO procedure [30,31]. First, viable vs nonviable cells were distinguished by washing the cells twice with phosphate-buffered saline (PBS), followed by incubation with Ghost Dye Red 780 (Cat. 13-0865-T100, Tonbo Bioscience) for 30 minutes at 2°C. The cells were subsequently washed twice with FACS buffer consisting of 1X Dulbecco’s phosphate-buffered saline (PBS) (Cat. 21-0310CV, Corning, NY), 1% bovine serum albumin (Cat. BP1600-100, Fisher Scientific, NH), and 0.1% sodium azide (Cat. S2002, Millipore Sigma, MA). Next, intracellular cytokine quantification was performed using the Transcription Factor Staining Buffer Kit (Cat. TNB-0607-KIT, Tonbo Biosciences, CA) following the manufacturer’s instructions. The cells were subsequently resuspended in Transcription Factor Fix/Permeabilization Buffer and incubated in the dark at 4°C for 1 hour. The samples were then washed with flow cytometry permeabilization buffer at 527 × g for 10 minutes at 4°C. The supernatant was discarded, and a mouse FcR blocker (Cat. 130-092-575, Miltenyi, GER) was added to the suspension, which was subsequently incubated for 5 minutes at room temperature. Two percent rat serum (Cat. 1355, Stemcell, CAN) was added and the mixture was incubated for an additional 15 minutes to reduce non-specific antibody binding. The following fluorochrome-conjugated monoclonal antibodies were added to the samples: anti-mouse IFN-γ violetFluor450 (Cat. 75-7311-U100, Tonbo Biosciences, CA), anti-mouse CD3ε BV711 (Cat. 100348, BD Biosciences, NJ), anti-mouse CD11c FITC (Cat. 35-0114-U025, Tonbo Biosciences, CA), anti-mouse CD19 PE (Cat. 50-0193-U100, Tonbo Biosciences, CA), anti-mouse CD8a PE-Cy5 (Cat. 55-0081-U25, Tonbo Biosciences, CA), anti-mouse CD25 PE-Cy7 (Cat. 60-025-U100, Tonbo Biosciences, CA), anti-mouse CD14 APC (Cat. 123312, BioLegend, CA), anti-mouse CD4 redFluor 710 (Cat. 80-0042-U100, Tonbo Biosciences, CA), anti-mouse CD69 APC (Cat. 20-0691-U025, Tonbo Biosciences, CA), anti-mouse IL-4 PE (Cat. 50- 7041-U025, Tonbo Biosciences, CA), and anti-mouse IL-17A FITC (Cat. 506907, BioLegend, CA). The samples were incubated in the dark for 30 minutes at room temperature, then washed with permeabilization buffer and centrifuged at 527 x g for 5 minutes. After the supernatant was discarded, the cells were fixed in 1% paraformaldehyde (Cat. 50-980-487, Fisher Scientific, NH). Data were acquired using an Aurora flow cytometer (Cytek, CA) and analyzed using Flow Jo software (v10) and a flow gating strategy designed to remove non-laminar artifacts (time vs. SSC-A), doublets (SSC-H vs SSC-A), debris (FSC-A vs SSC-A), and dead cells (GhostDye-780 vs SSC-A, Figure 1A).

**Figure 1:**
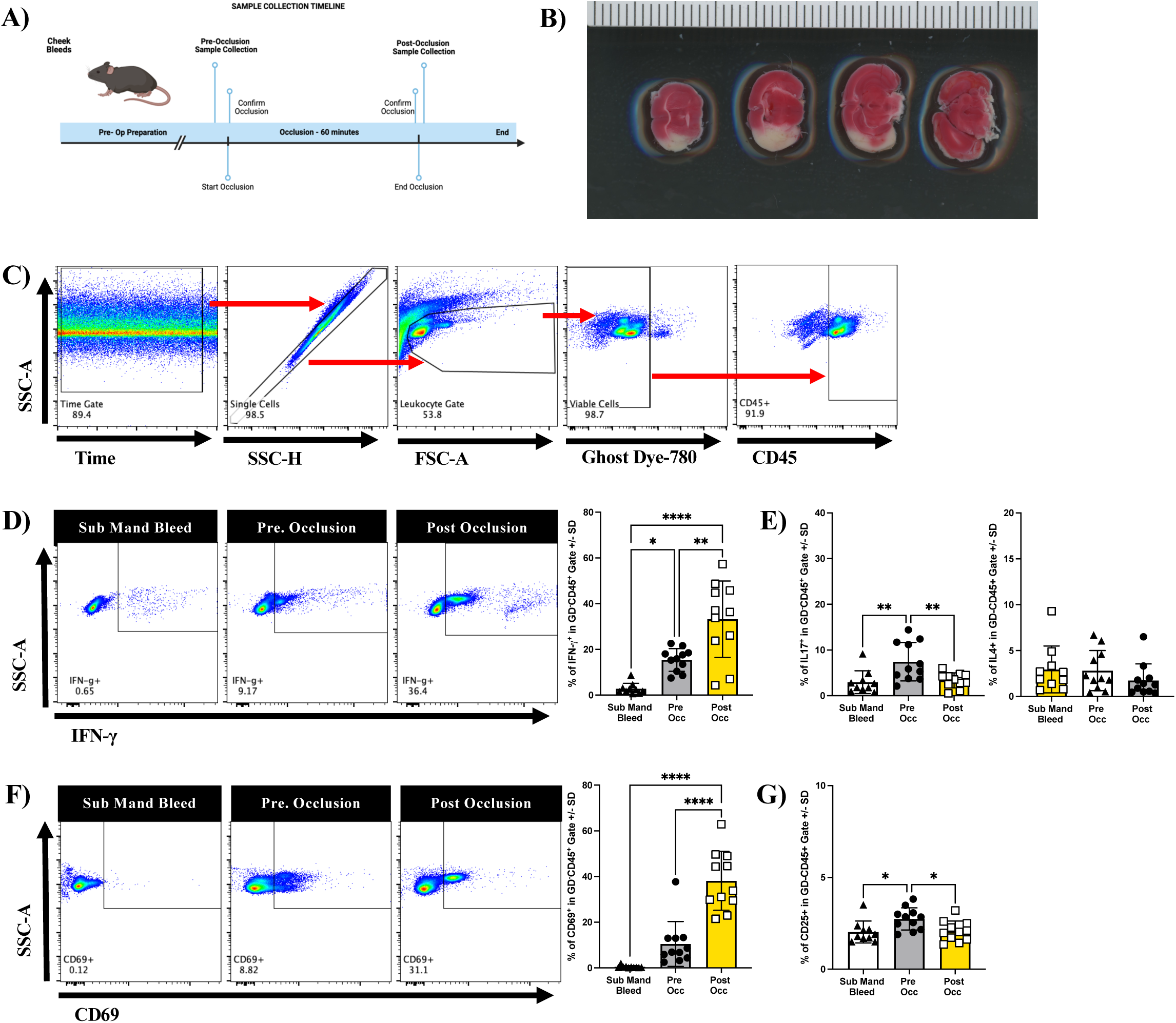
Novel inflammatory immune response immediately after experimental stroke. **A**) Schematic of the experimental timeline. The mice underwent 60-minute transient middle cerebral artery occlusion (tMCAO). Intra-arterial blood samples were collected pre-occlusion (prior to filament insertion) and post- occlusion (immediately after reperfusion). Submandibular bleeds from naïve mice served as distal, nonsurgical controls. **B**) Representative TTC-stained coronal brain slices confirming infarction after tMCAO. **C**) Flow cytometry gating strategy; exclusion of non-laminar flow events, doublets, and dead cells, followed by gating on CD45^+^ leukocytes from live single cells. **D**) Representative flow cytometry plots and cumulative data showing IFN-γ^+^ cells in the GD780^-^CD45^+^ leukocyte population across the indicated conditions. IFN-γ^+^ cells significantly increased in both pre-occlusion and post-occlusion samples. **E**) Quantification of IL-17^+^ and IL- 4^+^ cells among GD780^-^CD45^+^. IL-17^+^ cells were elevated in pre-occlusion samples only; IL-4^+^ cells showed no significant changes. **F**) Representative plots and cumulative data showing CD69^+^ cell frequency in GD780^-^ CD45^+^ leukocytes across indicated conditions. **G**) Quantification of CD25^+^cells, which were elevated only in pre-occlusion samples. The data are shown as the means ± SD. Each dot represents an individual animal (n=8– 10/group). Statistical analysis was performed using one-way ANOVA with Tukey’s post hoc test. *p< 0.05, ** p< 0.01, ****p< 0.0001.

### Mixed Cortical Culture

To study neuroimmune interactions *in vitro*, mixed cortical cultures (MCC) were generated as primary cell cultures containing a heterogeneous population of brain cells, including neurons, astrocytes, oligodendrocytes, and microglia [32–34]. Cortices were harvested from postnatal day 0–2 C57BL/6J pups bred in-house. The tissue was mechanically dissociated and incubated in 0.25% trypsin (Cat. 25200056, Fisher Scientific, NH) at 37°C for 15 minutes to facilitate enzymatic digestion. Following digestion, the mixture was suspended in an equal volume of MCC medium consisting of 1X Dulbecco’s modified Eagle medium (DMEM; Cat. MT15017CM, Fisher Scientific, NH), 10% fetal bovine serum (FBS; Cat.100-106-500, Gemini Bio, CA), 1% sodium pyruvate (Cat. 11360070, Fisher Scientific, NH), and 100 µg/mL Primocin (Cat. NC9141851, Fisher Scientific, NH). The suspension was subsequently centrifuged at 527 × g for 10 minutes at 4°C. The supernatant was discarded, and the cell pellet was resuspended in fresh MCC media and passed through a 40 µm nylon filter. The filtered suspension was then centrifuged again under the same conditions, and the resulting pellet was resuspended in fresh MCC media. Cell viability and counts were assessed using 4% trypan blue (Cat. 1525006, Gibco, NY) and a Cytosmart cell counter (Corning, NY). The cells were plated at a density of 2 × 10⁶ cells per well in Primaria six-well cell culture plates (Cat. 353846, Fisher Scientific, NH) [34]. MCC plates were placed in a humidified incubator at 37°C with 5% CO_2_. After three days, the cells were gently washed with fresh PBS and supplied with fresh MCC medium for an additional seven days. On day 10, the medium was replaced once more, and cultures were returned to the incubator until day 14, at which point they were used for downstream assays.

### Oxygen-Glucose Deprivation Model

To induce *in vitro* ischemia, mixed cortical culture (MCC) plates were subjected to oxygen–glucose deprivation (OGD). Cultures were washed three times with OGD media consisting of 1X DMEM without glucose (Cat. 11966-025, Gibco, NY) supplemented with 100 μg/mL Primocin. After washing, 2 ml of OGD medium was added to each well. The plates were then placed in a hypoxia chamber (Coy Laboratory Products, MI) regulated to 0.1% oxygen and incubated under hypoxic conditions for 2 hours and 30 minutes. Following the hypoxia exposure, the plates were removed from the chamber, and the plates were randomly divided into experimental groups for downstream analysis. As a normoxic control, parallel MCC plates were maintained outside the chamber in glucose-containing DMEM. All cultures (OGD and normoxic conditions) were subsequently co-cultured under standard incubator conditions. At the end of the incubation period, the samples were harvested for downstream analysis.

### Immunofluorescence Imaging

To analyze CNS cell populations in the *in vitro* experiments, immunofluorescence imaging was performed. After the experimental conditions, the wells were washed three times with PBS, followed by fixation with 4% paraformaldehyde (PFA, Cat. 50-980-487, Fisher Scientific, NH) for 15 minutes at room temperature. After fixation, the cells were washed with PBS and blocked for 60 minutes at room temperature with a blocking solution composed of 1X PBS, 10% goat serum (Cat, 100-109- 500, Gemini Bio, CA), and 0.1% Triton X-100 (Cat. 1610407, Bio-Rad Laboratories, CA). Afterwards, the mixture was aspirated, and the cells were incubated with an anti-mouse NeuN primary antibody (Cat. MAB377, Millipore Sigma, MA) at 4°C for 24 hours. After incubation, the cells were washed with PBS and then incubated with a goat anti-mouse IgG Alexa Fluor 594-conjugated secondary antibody (Cat. A11032, Invitrogen, MA) for 1 hour at room temperature in the dark. The cells were washed and counterstained with 0.2% DAPI (Cat. D9542, Millipore Sigma, MA) for 5 minutes at room temperature to visualize the nuclei.

Finally, the cells were washed, and a glass coverslip was mounted with Aqua-Poly/Mount. Blinded to experimental conditions, the investigator used the BZ-X810 fluorescence microscope (Keyence, IL) to capture fluorescence data. Four randomly selected fields per well were imaged, and cell counts were quantified using the BZ-X800 Analyzer Software (Keyence, IL).

### Statistics

Data are presented as the mean ± standard deviation (SD). Statistical analysis was performed using either a one-way or two-way analysis of variance (ANOVA), followed by Tukey’s post hoc test for multiple group comparisons, or an unpaired two-tailed Student’s t test for comparisons between two independent groups. Normality was assessed using the Shapiro–Wilk test prior to parametric testing. For data that did not meet the assumptions of normality, nonparametric alternatives (e.g., Kruskal-Wallis test or Mann-Whitney U test) were applied as appropriate. All analyses were performed using Prism v10 (GraphPad Software, CA). Experimental replicates (n) represent independent biological samples (e.g., individual animals or culture wells), as indicated in the figure legends. All experiments were independently replicated at least twice unless otherwise noted.

## RESULTS

### Inflammatory Cells Immediately after Experimental Stroke

Our previous clinical study revealed elevated levels of interferon-gamma (IFN-γ) in the post-ictus blood samples collected within 7 hours of stroke onset [33]. To bridge clinical findings with mechanistic insight, we investigated early cytokine responses following experimental stroke using a murine model. To determine whether this hyperacute pro-inflammatory signature could be recapitulated in a preclinical model, we employed a 60-minute transient middle cerebral artery occlusion model in C57BL/6J mice. Intra-arterial blood was collected immediately before filament insertion (pre-occlusion), followed by occlusion, and a second sample was collected immediately after reperfusion (post-occlusion) as outlined (**Figure 1A**). As a distal, nonsurgical control, peripheral blood was obtained by submandibular blood draw from naïve mice. Confirmation of successful occlusion was evaluated by cerebral blood flow reduction at the time of tMCAO induction, while confirmation of neuropathology was performed on brain slices using the TTC assay (representative image, **Figure 1B**).

Immediately after tMCAO induction, blood samples were processed for flow cytometry, and a gating strategy was applied to exclude nonlaminar flow artifacts, doublets, dead cells, and gated on CD45^+^ leukocytes (**Figure 1C**). Our prior study revealed changes in the quantities of inflammatory (IFN-γ and interleukin-17) and anti-inflammatory (interleukin-4) cytokines. Thus, in this study, we took the next step and sought to identify the source of these cytokines by quantifying IFN-γ-, IL-17-, and IL-4-producing immune cells. Compared to the control submandibular samples (2.791% ± 2.26), we observed a significant increase in GD780^-^CD45^+^IFN-γ^+^ cells in the pre-occlusion samples (15.41% ± 4.76, *p*<0.05) and a more robust increase in the post-occlusion samples (33.20% ± 15.94, *p*<0.000, **Figure 1D**). This increase was specific to IFN-γ- producing cells, as the IL-17^+^ cells were elevated only in pre-occlusion samples (7.44% ± 4.02, *p*<0.01) whereas the post-occlusion samples (3.36% ± 1.55) were comparable to the controls (2.95% ± 2.38, n.s., **Figure 1E**). IL-4^+^ cells showed no significant differences across all conditions (**Figure 1E**).

To determine whether the increase in IFN-γ^+^ cells was associated with standard activation markers, we quantified CD69^+^ (early activation marker) and CD25^+^ (late activation marker) cells [35,36]. Compared with those in the submandibular control samples, CD69^+^ cells trended to an increase in the pre-occlusion (10.48% ± 9.36, *p*=0.05) but significantly increased in the post-occlusion samples (38.12% ± 12.29, p<0.001, **Figure 1F**). This finding was specific to CD69, as CD25^+^ cells were found to be increased only in the pre-occlusion sample (2.73% ± 0.52, *p*<0.05, **Figure 1G**).

In summary, our data demonstrate a robust hyperacute increase in IFN-γ-producing CD45^+^ cells following experimental stroke, accompanied by early activation marker expression. This observation supports the novel hypothesis that stroke rapidly and immediately induces the activation of immune cells downstream of the ictus.

### CD14^+^ cells, a novel cellular source of IFN-γ in hyperacute stroke

Given the marked increase in IFN-γ^+^ cells observed in the post-occlusion samples, we next sought to identify its cellular source. To this end, we adjusted our flow cytometry panel to include key immune cell markers present in peripheral blood (**Supplementary Figure 1**). We first examined T-cells, which are conventionally associated with IFN-γ production. Analysis of the CD4 and CD8 T-cell populations revealed no significant difference in the percentage of IFN-γ-producing cells or a consistent relationship between pre- and post-occlusion samples (**Figure 2A**), indicating that these adaptive T-cell subsets are unlikely to account for the observed increase in IFN-γ during the hyperacute phase. Although significantly less abundant (2–5%) in murine blood, natural killer (NK) cells are also known producers of IFN-γ [37]. However, similar to the findings in the T-cell populations, we did not detect a significant change in IFN-γ^+^NK1.1^+^ cells between the pre- and post-occlusion samples (**Supplementary Figure 2**).

**Figure 2:**
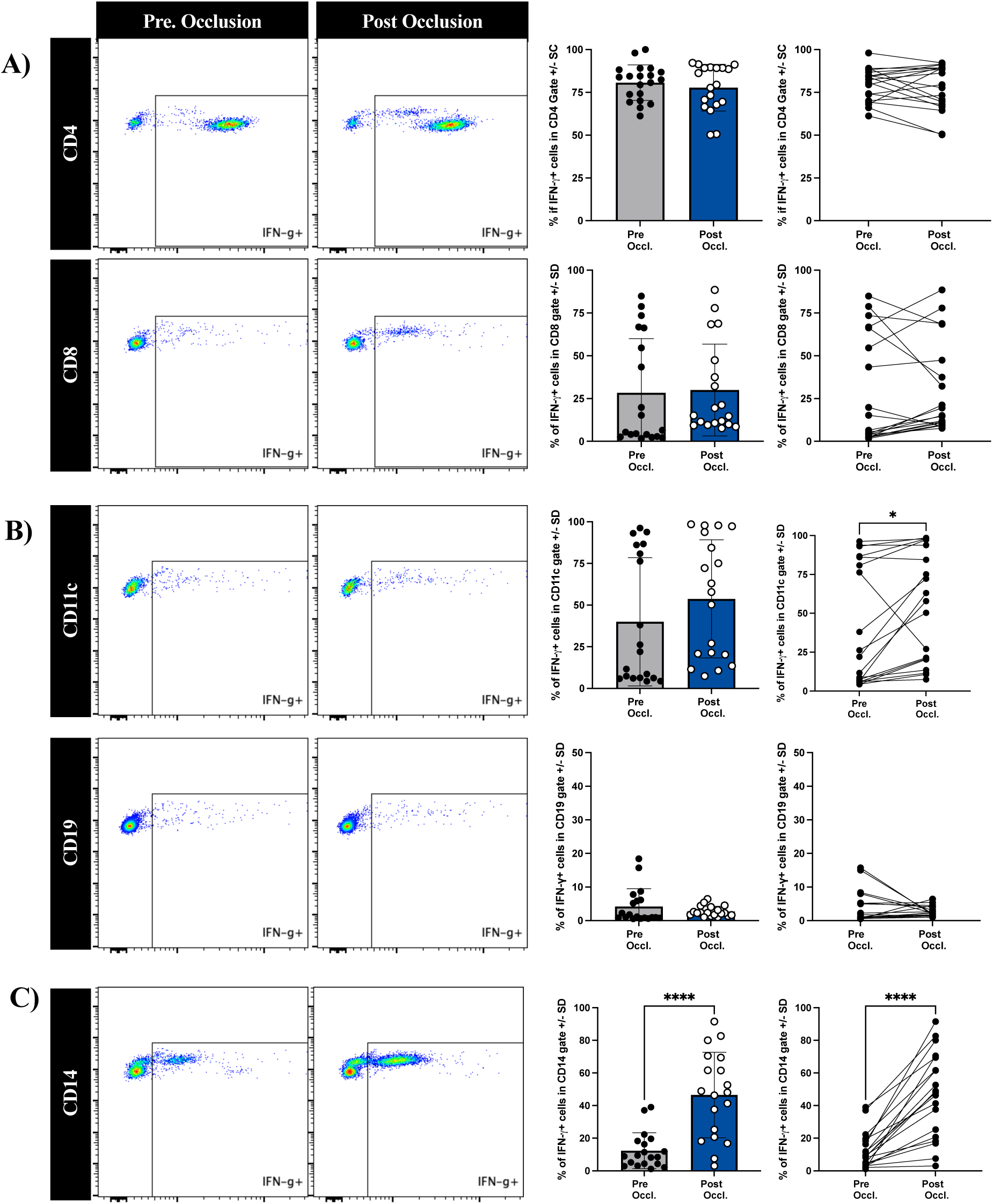
CD14^+^ cells are the predominant source of IFN-γ following experimental stroke. **A**) Representative flow cytometry plots (left) and quantification (center and right) of IFN-γ^+^ cells within CD4^+^ and CD8^+^ T-cell populations from pre- and post-occlusion arterial blood samples. No significant differences were observed between conditions, and paired analysis revealed no consistent relationship, suggesting CD4^+^ and CD8^+^ T-cells are not the primary contributors to increased IFN-γ levels during the hyperacute phase. **B**) IFN-γ expression in non-classical IFN-γ producing cell subsets, including CD11c^+^ dendritic cells and CD19^+^ B-cells. Quantification revealed no significant changes in the percentage of IFN-γ^+^ cells in either population. **C**) In contrast, IFN-γ^+^ cells within the CD14^+^ population were significantly increased in post-occlusion relative to pre-occlusion samples. Paired animal analysis confirmed this consistent increase across samples, identifying CD14^+^ monocytes as a major source of IFN-γ in the immediate post-ictus period. The data are presented as the means ± SD. Each dot represents an individual animal (n=19/group). Significance was determined by paired or unpaired Student’s t test as appropriate. *p< 0.05, ****p< 0.0001.

We then examined non-conventional sources of IFN-γ, including CD11c^+^ dendritic cells [38], CD19^+^ B cells [39], and CD14^+^ monocytes [24]. No differences were observed in the proportion of IFN-γ-producing cells within the CD11c^+^ or CD19^+^ populations between pre- and post-occlusion samples (**Figure 2B**), suggesting that B cells do not contribute substantially to the increase in hyperacute IFN-γ production, whereas the paired analysis revealed a small but significant increase in the percentage of CD11c^+^ cells (40.00% ± 38.49 vs. 53.72% ± 35.44, *p*=0.02). In contrast, compared with the pre-occlusion sample (12.31% ± 10.67), we observed a robust and significant increase in the percentage of IFN-γ-producing CD14^+^ cells in post-occlusion samples (46.48% ± 25.51, p<0.0001), with a consistent increase across paired animals (**Figure 2C**).

Together, these data indicate that the marked increase in IFN-γ-producing cells observed after tMCAO is driven primarily by CD14^+^ macrophages, rather than adaptive T or B-cell subsets. These findings support a previously unrecognized role for innate myeloid cells as early sources of IFN-γ following ischemic stroke.

### IFN-γ is differentially expressed in hyperacute stroke

To better characterize the cellular source of IFN-γ during the hyperacute phase of stroke, we stratified IFN-γ^+^ cells on the basis of their IFN-γ expression intensity and identified two distinct populations: a low-expression subset (IFN-γ^low^) and a high-expression subset (IFN-γ^hi^, **Figure 3A**). In the pre-occlusion samples, the relative frequencies of IFN-γ^low^ and IFN-γ^hi^ cells were comparable. However, post-occlusion samples demonstrated a significant expansion of the IFN-γ^low^ population (30.56% ± 10.30) compared with the IFN-γ^hi^ population (12.01% ± 4.04, p<0.001), indicating a shift in IFN-γ dynamics immediately following ischemic injury.

**Figure 3:**
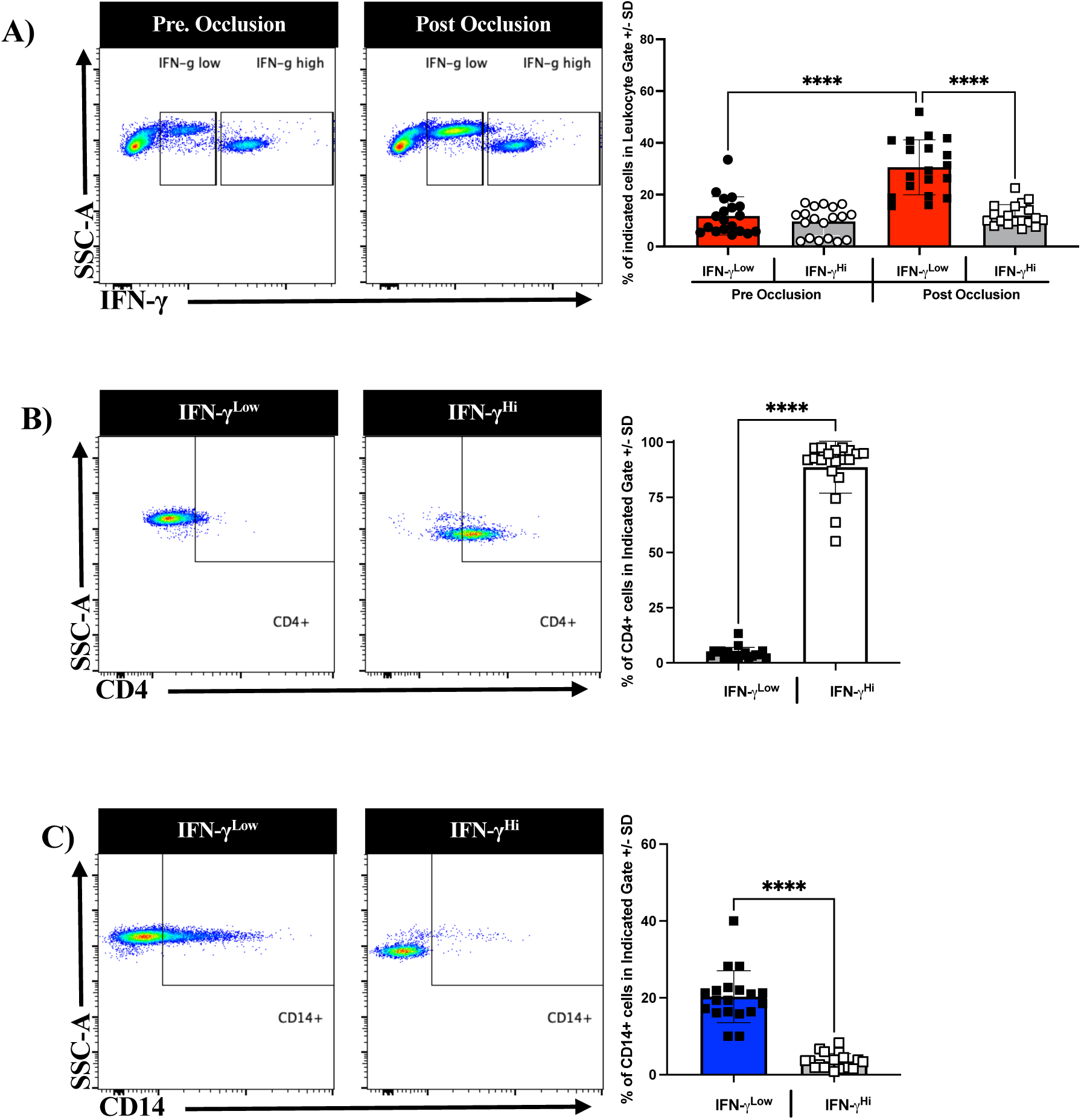
Stratification of IFN-γ-producing cells reveals distinct subsets with divergent cellular identities. **A**) Representative flow cytometry plot (left) showing gating of IFN-γ^low^ and IFN-γ^hi^ populations within the leukocyte populations from pre- and post-occlusion arterial blood samples. Quantification (right) revealed a significant expansion of IFN-γ^low^ populations in the post-occlusion sample relative to the IFN-γ^hi^ cells, indicating a shift in the nature of IFN-γ-producing cells following ischemic injury. **B**) Representative plots and quantification of CD4 expression within the IFN-γ^low^ and IFN-γ^hi^ gates. The IFN-γ^hi^ population was composed predominantly of CD4^+^ T-cells, which is consistent with conventional Th1-driven IFN-γ production. **C**) Representative plots and quantification of CD14^+^ cells within the IFN-γ^low^ and IFN-γ^hi^ gates. The IFN-γ^low^ population was enriched for CD14^+^ monocytes, indicating a previously underappreciated role for innate myeloid cells as early as IFN-γ producers following stroke. The data are presented as the means ± SD. Each dot represents an individual animal. Statistical significance was determined by an unpaired Student’s t-test. ****p< 0.0001.

To determine the cellular composition of these two IFN-γ-producing subsets within the post-occlusion samples, we stratified each population by lineage markers. Within the IFN-γ^hi^ cluster, CD4 T-cells were the predominant cell type (**Figure 3B**), which is consistent with the classical Th-1-mediated cytokine production [40]. In contrast, the IFN-γ^low^ cluster was composed primarily of CD14^+^ monocytes (**Figure 3C**). These findings suggest that monocytes serve as a distinct, non-classical source of IFN-γ in the hyperacute phase of stroke.

### Oxygen-Glucose Deprivation-mediated Injury is Sufficient to Induce IFN-γ Production by CD14^+^ cells

To further confirm that CD14^+^ cells are a source of IFN-γ during the hyperacute phase of stroke, we sought to identify the upstream insult capable of initiating this cytokine response. Using an established *in vitro* model of ischemia – oxygen-glucose deprivation (OGD), we simulated ischemic injury in CNS cells and assessed their ability to induce IFN-γ production in CD14^+^ cells by flow cytometry (**Figure 4A**). When naïve splenocytes were co-cultured with OGD-treated CNS cells, no significant increase in IFN-γ^+^CD14^+^ cells was observed compared with that in normoxia-treated controls (**Figure 4B**). Likewise, treatment of splenocytes with supernatant from OGD-treated CNS cultures failed to elicit a significant production of IFN-γ from CD14^+^ cells. In contrast, the combination of both OGD-treated CNS cells and their supernatants induced a significant increase in IFN-γ^+^CD14^+^ cells, relative to their normoxia-treated controls (31.82% ± 8.66 vs. 51.53% ± 2.25, *p*<0.05), which was also significant compared to the singular supernatant or OGD-treated CNS conditions (31.64 ± 11.49, p<0.05, or 30.12 ± 17.44, *p* <0.01, respectively, **Figure 4B**). Taken together, these data suggest that both soluble and cell-associated signals act synergistically to trigger this response.

**Figure 4:**
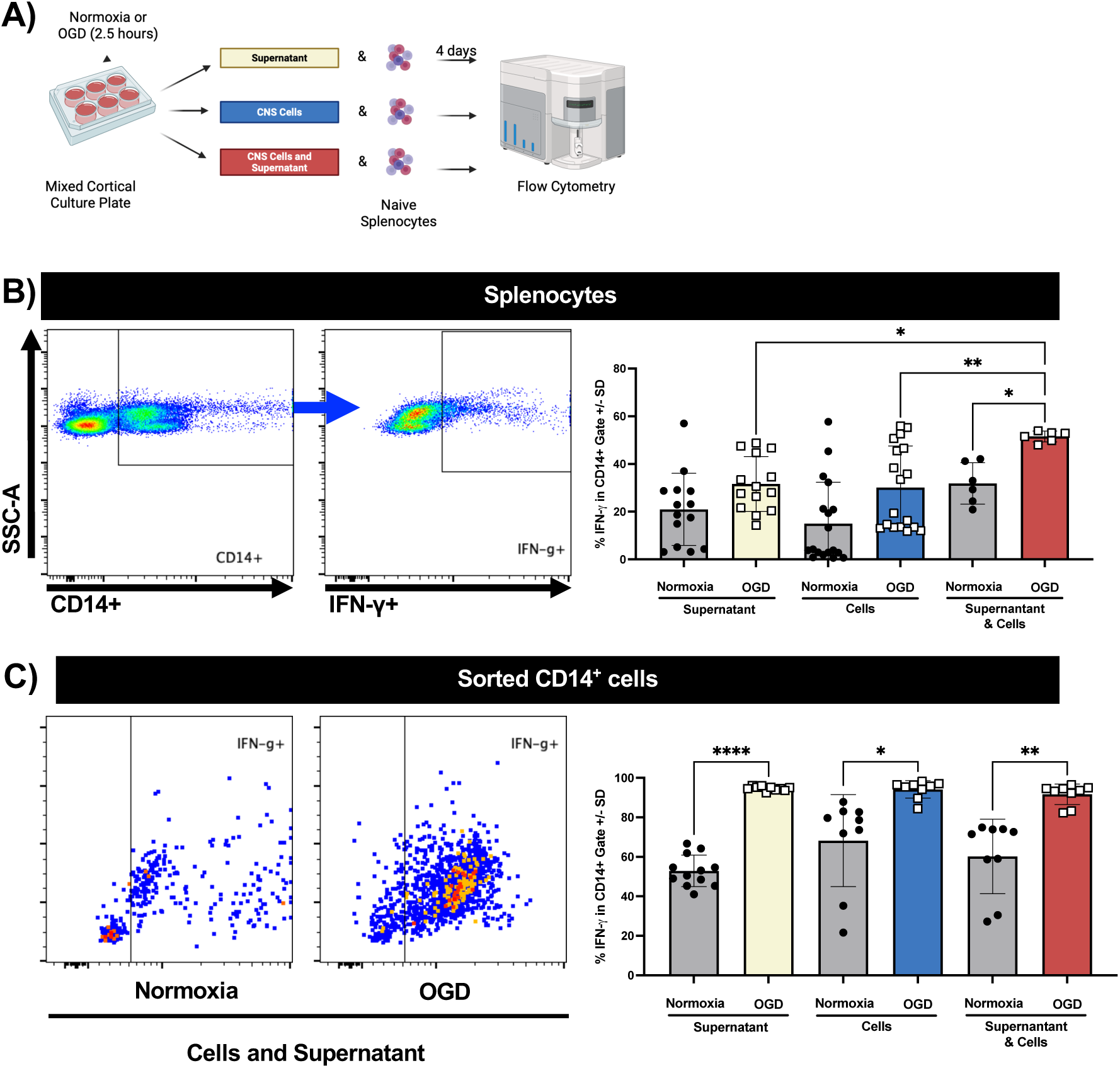
Oxygen–glucose deprivation (OGD)-injured CNS cells and their supernatants stimulate IFN- γ production in CD14^+^ cells. **A**) Schematic representation of the in vitro ischemia model. Mixed murine cortical cultures were subjected to normoxia or oxygen–glucose deprivation (OGD) for 2.5 hours. After treatment, CNS cells, their supernatants, or both were co-cultured with naïve splenocytes for four days, followed by flow cytometry analysis. **B**) Representative flow cytometry plots (left) and quantification (right of IFN-γ^+^ cells within CD14^+^ populations in naïve splenocytes post-culture. Compared with normoxic controls, neither OGD-treated CNS cells nor their supernatants alone significantly increased IFN-γ production. However, the combination of OGD-treated CNS cells and supernatants induced a significant increase in IFN- γ^+^CD14^+^ cells, suggesting a synergistic effect of soluble and cell-associated signals from ischemic CNS cells. **C**) Compared with their respective normoxic controls, sorted CD14^+^ cells stimulated under the same conditions presented robust and significant increase in IFN-γ^+^ cells across all OGD-treated. This indicates that both secreted and contact-dependent injury signals are sufficient to directly activate CD14^+^ cells in the absence of other immune cell types. The data are shown as the mean ± SD. Each dot represents an individual biological replicate. Statistical significance determined by one-way ANOVA with Tukey’s post hoc test. *p< 0.05, ** p< 0.01, ****p< 0.0001.

To determine whether these injury-associated factors directly stimulate CD14^+^ cells, we isolated CD14^+^ cells from naïve splenocytes and exposed them to the same conditions. Interestingly, purified CD14^+^ cells responded robustly to each of the three conditions – OGD-treated supernatant, OGD-treated CNS cells, and the combined condition, with significant increases in IFN-γ^+^CD14^+^ cells observed in all the cases (**Figure 4C**). These findings indicate that both secreted and contact-dependent signals from ischemia-injured CNS cells are sufficient to directly activate CD14^+^ cells and promote IFN-γ production in the absence of other immune cells.

### Oxygen and Glucose Deprivation of CNS Cells Primes Splenocytes to Exacerbate Neuronal Injury

To investigate whether ischemia-induced immune responses are associated with cytotoxic effects on CNS cells, we treated CNS cells (MCC plates) with OGD and co-cultured them with naïve splenocytes. Using immunofluorescence, we quantified the total number of CNS cells (DAPI, blue) and neurons (NeuN, red, **Figure 5A**). Compared with the normoxic controls, OGD treatment alone resulted in a substantial reduction of total CNS cells (DAPI^+^) in the MCC plates (2447 ± 299 vs 861.4 ± 291, *p*<0.0001, **Figure 5B).** The addition of splenocytes to OGD-treated CNS cells further reduced DAPI^+^ counts relative to OGD-alone (424.8 ± 285, *p*<0.0001), suggesting that peripheral immune cells exacerbate ischemia-induced cell loss. Interestingly, when OGD-treated cells were co-cultured with splenocytes and their corresponding OGD-derived supernatants, the DAPI^+^ cell counts were greater than the splenocytes + OGD-treated CNS cell condition (1085.9 ± 451.3, *p*<0.0001), suggesting an alteration of the immune-CNS cell interactions driven by soluble inflammatory factors.

**Figure 5:**
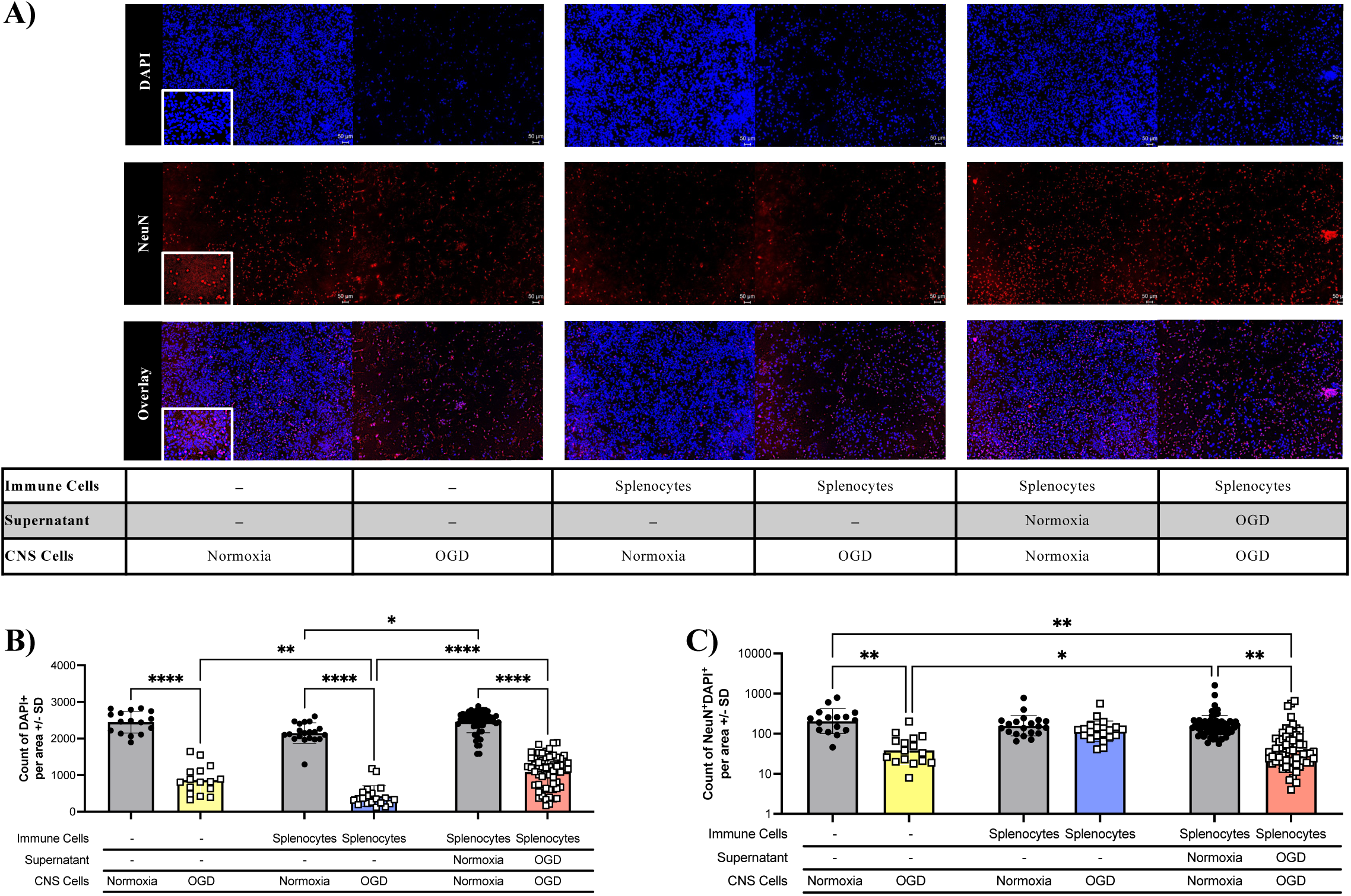
Co-culture of splenocytes with OGD-injured CNS cells exacerbates neuronal loss. **A**) Representative immunofluorescence images of DAPI+ nuclei (blue) and NeuN^+^ neurons (red) from CNS cultures subjected to normoxia or oxygen–glucose deprivation (OGD), with and without naïve splenocytes co-culture and/or CNS-injured supernatant. The white insets show magnified regions. **B**) Quantification of the total cell count (DAPI^+^ nuclei per area ± SD across experimental conditions). Compared with normoxia, OGD alone significantly reduced total cell numbers. Co-culture of splenocytes with OGD-treated CNS cells further decreased DAPI^+^ counts, whereas addition of OGD-derived supernatant to OGD-treated CNS cultures with splenocytes resulted in cell counts similar to OGD-treated CNS cells alone. **C**) Quantification of NeuN^+^DAPI^+^ neuronal counts per area ± SD. OGD treatment significantly reduced neuronal survival, and loss was further exacerbated when splenocytes were co-cultured with OGD-treated CNS cells and their supernatant, corresponding to a condition that also induced a marked increase in IFN-γ^+^CD14^+^ cells. The data are shown as the means ± SD. Statistical significance was determined by one-way ANOVA with Tukey’s post hoc test. Each dot represents an independent biological replicate. Scale bar = 50 μm. *p< 0.05, ** p< 0.01, ****p< 0.0001.

To evaluate the impact on neurons specifically, NeuN^+^DAPI^+^ cells were quantified. Compared with normoxic controls, OGD alone significantly decreased neuronal counts (258 ± 194 vs. 53 ± 48.35, *p*<0.01, **Figure 5C**). This loss of neurons was reversed when splenocytes were added (139 ± 113.5, n.s.). Taken together with the observation that this condition resulted in a significant reduction in total CNS cells (**Figure 5B**), we speculate that the addition of splenocytes to OGD-induced CNS cells results in cytotoxicity to non-neuronal cell populations. In contrast, when OGD-derived supernatant was added to the OGD-treated CNS cells and splenocytes, we observed a reduction of NeuN^+^DAPI^+^ cells (198.8 ± 224.4 vs. 75.52 ± 124.5, *p*<0.01). Concomitant with the increase in total CNS cells (**Figure 5B**) and the significant increase in IFN-g^+^CD14^+^ cells (**Figure 4B**), these findings suggest that ischemic injury to CNS cells causes a significant increase in inflammatory IFN-g^+^CD14^+^ cells that is associated with neuronal loss.

## DISCUSSION

Stroke initiates a rapid and complex immune response that evolves and contributes significantly to both acute injury and long-term neurological outcomes. While much attention has been given to subacute and chronic post-stroke inflammation, the hyperacute phase (within hours of ischemia onset) remains comparatively understudied. In addition, most work on stroke immunology uses venous blood, so analysis of the arterial compartment remains largely unexplored and underappreciated. Our prior clinical investigation identified elevated levels of IFN-γ in the downstream arterial vascular compartment of patients within seven hours of large-vessel occlusion, indicating early immune activation at the site of ischemia [33]. The present study builds upon these findings by using a murine stroke (tMCAO) model to dissect the cellular source, stimuli, and potential impact of IFN-γ production in the hyperacute post-stroke environment.

Consistent with our human findings, the percentage of IFN-γ-producing cells were significantly elevated in post-occlusion arterial blood collected within 1-2 hours of ischemia. This early increase occurred in the absence of IFN-γ elevation from canonical sources (T- or NK-cells), indicating instead that intra-arterial CD14^+^monocytes are a novel and dominant early source. IFN-γ is a cytokine that plays a critical role in immune responses [41] and serves as a key activator of macrophages [14]. It does so by overriding inhibitory feedback mechanisms [42] and driving macrophages into an activated state [43]. While IFN-γ is primarily produced by T-cells and natural killer cells [44], it can also be secreted by other cell types, including dendritic cells [22,45,46], B cells [47], macrophages [25], and microglia [48]. Pre-clinical studies have linked IFN-γ to worse stroke outcomes. Lambertsen *et al*. reported that transgenic mice expressing IFN-γ under the myelin basic protein promoter had larger infarct volumes compared to wild-type mice [49]. Similarly, Zhang *et al*. reported that IFN-γ-deficient mice subjected to tMCAO had reduced infarct volumes compared to wild-type controls three days post-stroke [50]. Additional studies have confirmed that reducing or eliminating IFN-γ leads to smaller infarct volumes, suggesting that IFN-γ contributes to worse stroke recovery outcomes [51,52]. This study identifies CD14^+^ cells, commonly known as monocytes/macrophages, as an unconventional sources of IFN-γ. Monocytes and macrophages are early responders after stroke, appearing within twenty-four hours and persisting for up to seven days within the ipsilateral hemisphere of an infarcted brain [53,54]. Monocyte levels are known to increase following stroke, with CD14^hi^CD16^+^ monocytes linked to worsening infarctions in stroke patients [55]. In-depth, preclinical studies suggest that they may play a pathogenic role in stroke progression. For example, in CCR2 knockout mice subjected to tMCAO, reduced monocyte infiltration was associated with small infarct volumes and improved blood–brain barrier integrity [56]. Similarly, in a clodronate depletion model, the macrophage-depleted tMCAO mice presented reduced demyelination and improved neurological outcomes [57]. These findings suggest that targeting monocytes/macrophages could represent a novel immunotherapeutic strategy to improve stroke recovery. However, the role of these cells in stroke remains complex. While they contribute to inflammation and injury early on, they may play a role in tissue repair during later phases [58,59]. This may explain why broad targeting of immune cells has already proven ineffective in stroke treatment [60]. In the present study, the CD14^+^ cells were isolated from the intra-arterial blood source; thus, we referred to these CD14^+^ cells specifically as monocytes.

Flow cytometry further revealed two distinct IFN-γ-producing populations: an IFN-γ^low^ population and an IFN-γ^hi^ population, of which the IFN-γ^low^ population increased post-occlusion. The IFN-γ^hi^ population was found to be composed primarily of CD4^+^ T-cells, whereas the IFN-γ^low^ population was largely composed of CD14^+^ monocytes. This is an unexpected finding given that monocytes are known to respond robustly to IFN- γ by regulating antigen presentation and enhancing antimicrobial and inflammatory functions. Direct monocyte-derived IFN-γ production has rarely been reported and has been described primarily under conditions of intense innate immune activation or severe inflammatory stimuli [61,62]. The relatively lower IFN-γ production (IFN-g^low^) by monocytes compared to T-cells may suggest a distinct biological role, perhaps modulating rather than purely exacerbating early neuroinflammation. The complexity of the role of IFN-γ in CNS inflammatory disease has also been extensively studied in multiple sclerosis (MS) and its animal model, experimental autoimmune encephalomyelitis (EAE). In these contexts, IFN-γ can act either as a pathogenic or a protective factor, depending on the timing, level, and source of production. Early administration or endogenous IFN-γ expression during EAE can paradoxically suppress disease by limiting T-cell expansion (immunoregulation) and promoting the apoptosis of autoreactive cells [16,18,31,63–65]. Conversely, elevated IFN-γ production during established or chronic phases of disease has been associated with increased demyelination, enhanced neuroinflammation, and exacerbation of clinical symptoms, underscoring its pathogenic potential [66–68]. Such dual and context-dependent roles for IFN-γ could similarly apply in ischemic stroke, where monocyte-derived IFN-γ may function with or distinct from classically lymphocyte- derived IFN-γ.

To investigate the upstream triggers of IFN-γ production, we employed an *in vitro* oxygen–glucose deprivation (OGD) model to simulate ischemic injury. While neither OGD-conditioned CNS cells nor their supernatants alone induced IFN-γ production, the combination of both significantly activated IFN-γ in CD14^+^ cells. These findings suggest that both cell-associated and soluble damage-associated molecular patterns (DAMPs) or cytokines are required to stimulate monocyte activation, which is consistent with the known synergy of IL-12 and IL-18 in IFN-γ production [69–72]. Notably, when CD14^+^ cells were isolated from splenocytes and cultured, we observed an enhanced IFN-γ production across all conditions, including OGD- conditioned supernatant alone, OGD-treated cells alone, and their combination (**Figure 4B**). This finding indicates that CD14^+^ cells are directly responsive to ischemia-associated stimuli and may not require accessory immune cells for activation, as previously assumed[73,74]. Rather than dampening activation, isolation may remove inhibitory regulatory inputs from other immune subsets, thus unmasking an intrinsic capacity of IFN- γ production in response to CNS injury. Notably, our *in vivo* studies detected IFN-γ production almost immediately after stroke, whereas our *in vitro* studies detected IFN-γ production at four days post-OGD. This delay may be due to limitations of the OGD model, which does not fully recapitulate the complex stroke environment, potentially leading to a slower response. In summary, these results suggest that CD14^+^ cells act as a highly sensitive immune sensors in the hyperacute phase of stroke and may respond directly to DAMPs released by ischemic CNS cells.

To assess potential effector functions, we co-cultured splenocytes with OGD-treated CNS cells and quantified both total CNS cell counts (DAPI^+^) and neurons (NeuN^+^DAPI^+^) in our mixed cortical culture plates (MCC). OGD-treated CNS cells alone resulted in a substantial reduction in both DAPI and NeuN cells, confirming that ischemia itself is sufficient to induce widespread CNS damage. Notably, the addition of splenocytes to OGD-treated CNS cells resulted in the greatest loss of total CNS cells, but neuronal counts were relatively preserved. These findings suggest that splenocytes exacerbate injury primarily in non-neuronal populations, such as astrocytes or oligodendrocytes.

Interestingly, when OGD-conditioned supernatant was added to these co-cultures, the number of neurons sharply decreased to levels comparable to those in the OGD alone group, whereas total cell loss was less severe than that in the OGD-CNS + splenocyte group. This implies that soluble ischemic factors enable or amplify neuron-specific injury, likely by unmasking cytotoxic pathways that are dormant in the presence of injured CNS cells and immune cells alone. Possible mechanisms include IFN-γ-driven modulation of glial support, enhanced oxidative stress, and death receptor pathway activation [75–78]. Alternatively, OGD supernatant may contain pro-inflammatory cytokines or DAMPs that prime immune cells for neurotoxic activity. Surprisingly, the addition of naïve splenocytes to OGD-treated CNS cells alone appeared to reverse neuronal loss observed with OGD injury alone. Despite inducing the most significant reduction in total DAPI^+^ CNS cells – suggesting enhanced glial or oligodendrocyte loss, NeuN^+^ neuronal counts were better preserved in this condition than with OGD alone. These findings suggest that naïve immune cells may exert a modulatory or even neuroprotective effect in the absence of inflammatory priming. One possibility is that resting splenocytes secrete regulatory cytokines, such as IL-10 or TGF-β, or support neuronal survival indirectly by clearing dysfunctional or apoptotic glia or debris [79–81]. However, when OGD-conditioned supernatant was introduced, this protective effect was abolished, and neuronal injury was restored to levels seen with OGD alone. This implies that soluble ischemic factors may reprogram immune cells towards a neuropathogenic phenotype, disrupting any protective role they might otherwise play. These context-dependent findings highlight the complex and bidirectional role of immune cells in early stroke and underscore the need to consider microenvironmental signals when assessing immune-mediated injury.

Despite evidence of IFN-γ production and CNS cell injury, the absence of overt neurotoxicity under some co-culture conditions suggests that the timing and maturation state of CD14^+^ cells may be critical. Our data revealed an increase in CD69^+^ and a lack of CD25^+^ expression, suggesting that these cells are still undergoing activation. This finding aligns with prior findings in EAE, where early IFN-γ can be regulatory, whereas delayed IFN-γ exacerbates disease [82].

These results highlight the dual potential role of CD14^+^ monocytes in stroke: as early amplifiers of inflammation and possible effectors of CNS injury. Importantly, this study was designed as a hypothesis- development study with the goal of replicating a human stroke phenotype in an animal model. Future studies are essential to better define the role of IFN-γ^+^CD14^+^ cells in stroke. For example, our current findings, which focused on CD14^+^ cells, can be improved by incorporating additional markers to distinguish these cells more clearly within the myeloid lineage, particularly macrophages. This can be achieved using classical macrophage markers such as F4/80 and CD68 [83]. Once confirmed as macrophages, further characterization can be performed using M1 markers, including CD80, CD86, and inducible nitric oxide synthase (iNOS), to refine the understanding of their phenotype and functional state [84]. Given the failure of global immunosuppression in stroke [85,86], targeting this early, IFN-γ-producing monocyte subset offers a novel and focused therapeutic opportunity.

The discovery of IFN-γ-producing CD14^+^ cells in the hyperacute phase of stroke challenges the traditional understanding of monocyte function and reveals a potentially novel mechanism of immune activation during this critical window. It must be highly noted that this study has identified a rare population of IFN-γ^+^CD14^+^ cells in the peripheral arterial blood, suggestive of monocytes with non-classical cytokine expression. While monocytes are not typical sources of IFN-γ, this phenotype may reflect an atypical activation state, which warrants further validation. In summary, the implications of this finding are significant as post-ictus IFN- γ^+^CD14^+^ cells can locally release cytokines that are known to disrupt the blood–brain barrier, amplify pro-inflammatory signaling, and recruit additional immune cells to the site of ischemia and potentiate ischemic injury. These activities may worsen acute tissue damage but could also influence subsequent long-term immune and repair responses. Further research into the origins, regulation, and functional impact of IFN-γ production by CD14^+^ cells after a stroke is essential.

## Conclusion

In summary, this study identifies CD14^+^ monocytes as an unrecognized and potent source of IFN-γ during the hyperacute phase of ischemic stroke, occurring within hours of vascular occlusion. Our *in vivo* and *in vitro* data demonstrate that these cells can be rapidly activated by both soluble and cell-associated factors released from ischemia-injured CNS tissue, leading to a pro-inflammatory state that coincides with early neuronal and non-neuronal cell loss. These findings extend current paradigms of stroke immunopathology by implicating innate myeloid cells – traditionally viewed as downstream effectors in the immediate cytokine milieu after stroke onset. The identification of this temporally and spatially restricted immune phenotype offers a unique opportunity for targeted intervention, where modulating IFN-γ-producing monocytes during the hyperacute window may limit secondary injury without impeding later reparative processes. Future work aimed at defining the molecular drivers, CNS targets, and long-term consequences of hyperacute IFN-γ production will be critical for translating these insights into clinically viable immunomodulatory strategies.

## Abbreviations

ACD: acid citrate dextrose
BBB: blood–brain barrier
CD: cluster of designation
CNS: central nervous system
DAMPs: damage-associated molecular patterns
DAPI: 4’6-diamidino-2-phenylindole
EAE: experimental autoimmune encephalomyelitis
IFN-γ: interferon gamma
IL-1: interleukin 1
IL-6: interleukin 6
IL-17: interleukin 17
iNOS: inducible nitric oxide synthase
LPS: lipopolysaccharide
MCC: mixed cortical culture
MMP9: matrix metallopeptidase 9
MS: multiple sclerosis
NK: natural killer cells
NeuN: neuronal nuclei
OGD: oxygen–glucose deprivation
PBS: phosphate-buffered saline
TGF-β: transforming growth factor beta
TNF-α: tumor necrosis factor alpha
tMCAO: transient middle cerebral occlusion
TNF-α: tumor necrosis factor alpha

## Acknowledgements

The data presented herein were obtained at the Flow Cytometry Facility, a core research facility at the University of North Texas Health Science Center, with the training and assistance of Dr. Sharad Shrestha. The image for Figures 4 was created using BioRender.

## Author Contributions

K.H., E.J.P., S.S., N.O., S.B.O., and N.J. performed the experiments. K.H. and S.B.O. designed the experiments, analyzed the data, and wrote the manuscript.

## Availability of Data and Materials

The datasets used and/or analyzed during the current study are available from the corresponding author upon reasonable request.

## Ethical Approval

The animal experiments were carried out in accordance with experimental protocols approved by the University of North Texas Health Science Center Institutional Animal Care and Use Committee (IACUC 2021-0045).

## Consent for publication

Not applicable

## SUPPLEMENTARY FIGURE

**Supplementary Figure 1: Flow cytometry gating strategy for identification of the IFN-γ-producing cell source.** Peripheral blood was immediately collected from the mice pre- and post-occlusion, processed for flow cytometry, and analyzed using a sequential gating approach. The cells were first gated on time vs SSC to exclude acquisition artifacts, followed by FSC-H vs FSC-A to isolate singlets. Viable leukocytes were identified based on the basis of FSC-A vs SSC-A characteristics. From the leukocyte gate, T-cells (CD3^+^) are further divided into CD4^+^ and CD8^+^ populations. CD3^-^ cells were gated to identify additional subsets: B-cells (CD19^+^), monocytes (CD14^+^), dendritic cells (CD11c^+^), and NK cells (NK1.1^+^). These populations were subsequently used to assess the proportion of IFN-γ^+^ cells within each subset in pre-and post-occlusion samples.

**Supplementary Figure 2: Natural killer (NK) cells do not contribute to the increase in IFN-γ production during the hyperacute phase of stroke.** Peripheral blood collected from mice pre- and post-occlusion was analyzed by flow cytometry to determine the proportion of IFN-γ-producing NK cells. NK cells were identified as CD3^-^NK1.1^+^ events within the leukocyte gate (see Supplementary Figure 1 for the gating strategy), and the percentage of IFN-γ cells within the NK1.1^+^ population was calculated. A representative flow dot plot of IFN- γ expression in NK1.1^+^ cells is shown for pre- and post-occlusion samples (left). Quantification (middle) and matched-pair analysis (right) revealed no significant differences in IFN-γ^+^ NK cells frequency between pre- and post-occlusion samples, suggesting that NK cells are not a major source of hyperacute IFN-γ after stroke.

