## Supplementary Figures for "Uncovering a new player in ischemic stroke: a study of intra-arterial interferon-gamma-producing monocytes in hyperacute stroke"

### Supplementary Figure 1: Gating Strategy for IFN-γ cell source

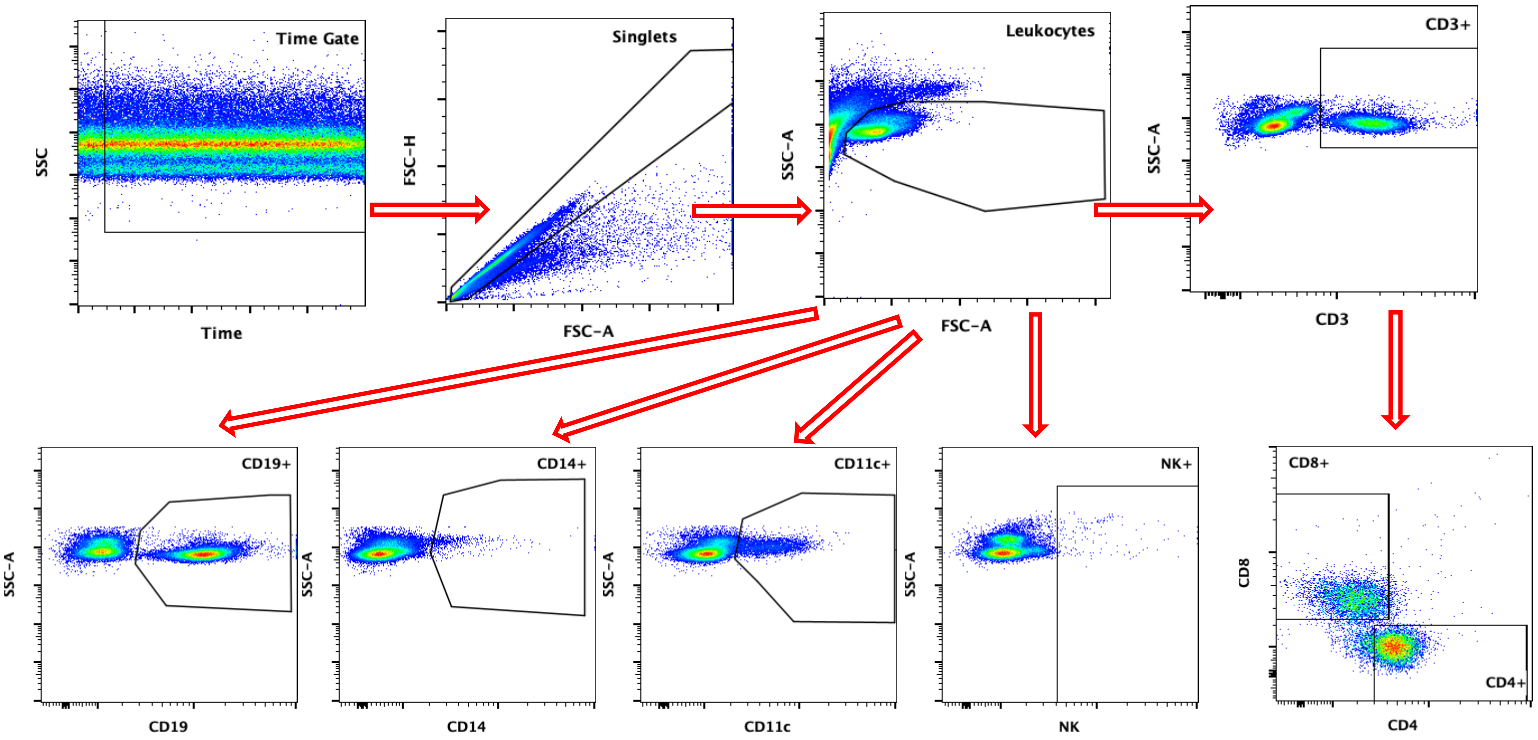

### Supplementary Figure 2: IFN-γ production by NK cells

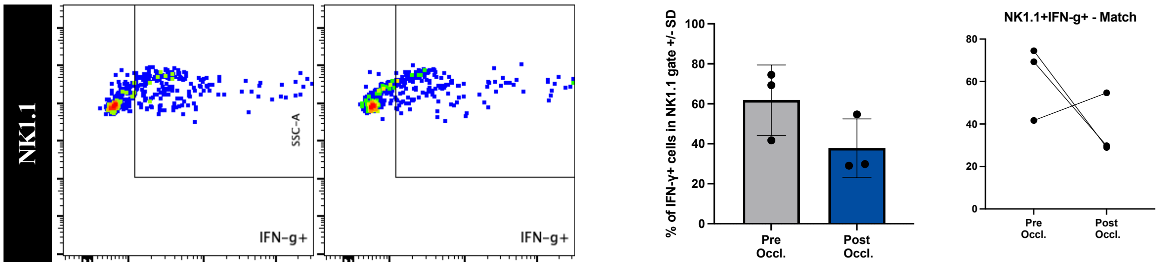
